# Mapping Lifespan Trajectories of Cognitive Flexibility with a Continuous Probabilistic Reversal Learning Measure

**DOI:** 10.64898/2026.06.18.733237

**Authors:** Mohammad Jowkar, Marjan Makhsous, Ehsan Rezayat

## Abstract

Cognitive flexibility is the ability to change the way of responding when the demands of the environment change. This study tested how cognitive flexibility develops across the lifespan. We used a new computerized task that gives a continuous score instead of just right or wrong answers. 221 healthy adults aged 18 to 71 completed the Continuous-score Probabilistic Reversal Learning Test (CPRLT). We calculated mean absolute error and adjusted error for rule-based learning, and fitted a Rescorla-Wagner model to estimate each person’s learning rate (alpha) for reward-based learning. All three scores have one breakpoint, performance improved rapidly from childhood to young adulthood, then declined slowly. Rule-based learning peaked around age 20. Reward-based learning peaked earlier, around age 18. This suggests that reward-based learning matures before rule-based learning. The pattern fits with brain development: reward circuits mature earlier, while prefrontal regions for rule-based learning develop later. Our continuous measure captured this difference, which binary tasks would miss.

## Introduction

Cognitive flexibility is the ability to adjust thoughts and behavior when environmental demands change. It enables learning, problem-solving, and goal-directed action. Without it, behavior becomes rigid and maladaptive (Hohl & Dolcos, 2024).

The neural basis of cognitive flexibility involves the prefrontal cortex and its connections with parietal and striatal regions (Uddin, 2021). These frontoparietal and frontostriatal networks undergo prolonged maturation during childhood and adolescence and show decline in older adulthood (Kupis & Uddin, 2023). Age-related changes in white matter integrity and cortical thickness within these networks are thought to contribute to behavioral inflexibility. This developmental trajectory extends across the lifespan, with performance improving through childhood and adolescence, peaking in young adulthood, and gradually declining thereafter (Tong et al., 2024).

Previous studies have mapped cognitive flexibility across the lifespan. Most have compared young adults with older adults. These studies consistently show that older adults perform worse than younger adults (Kupis & Uddin, 2023). A few studies have included participants across a wider age range. They found that flexibility improves from adolescence into early adulthood and then gradually declines (Cepeda et al., 2001; Huizinga et al., 2006). Some evidence suggests that peak performance occurs between the late teens and early thirties, depending on the specific task demands and the aspect of flexibility being measured (Tong et al., 2024). Other work has shown that rule-based learning and reward-based learning follow different developmental timecourses (Sandbrink & Summerfield, 2024). For example, reward-based learning may mature earlier, while rule-based learning continues to develop into early adulthood (Sandbrink & Summerfield, 2024). What is less clear is whether these two components peak at the same age or at different points. The present study addresses this question using a new continuous-error task that separates rule-based learning from reward-based free learning.

Another limitation is measurement. Traditional flexibility tasks, such as the Wisconsin Card Sorting Test (Berg, 1948)or task-switching paradigms (Ravizza & Carter, 2008), often produce binary outcomes – correct or incorrect. These measures do not capture the graded nature of adaptive learning. A participant may be close to the optimal response or far from it, but binary scoring treats both as errors. Recent work in working memory has shown that continuous measures, such as recall precision, reveal age-related changes that binary accuracy measures miss (Esfahan et al., 2025). The same logic applies to cognitive flexibility.

Therefore, we decided to use a new task, which gives continuous score in each trial, to study the development of cognitive flexibility through life span.

## Methodology

### Participants

221 participants (126 female) took part in the study. They were recruited through convenience sampling. To take part, people needed normal or corrected-to-normal eyesight and no diagnosed psychiatric disorder. We checked this by asking two simple questions: “Have you been diagnosed with any psychiatric disorder?” and “Do you have normal or corrected-to-normal eyesight?”. Before starting, all participants (or their legal guardians) read and signed a consent form. All procedures were approved by the Ethics Committee of the University of Tehran (IR.UT.PSYEDU.REC.1404.064) and were in accordance with the Declaration of Helsinki of 1964 and its later amendments.

### Measures

Continuous-score Probabilistic Reversal Learning Test (CPRLT): This test is taken from a study by Jowkar )2025). It has 150 trials. In each trial, a green circle appears on the screen (see Figure 1). The participant moves the mouse along a horizontal line to change the circle’s size, then clicks to choose a size (sizes are scaled from 0 to 1). After clicking, they get a score between -100 and 100. Their goal is to get as high a total score as possible.

**Figure 1.**
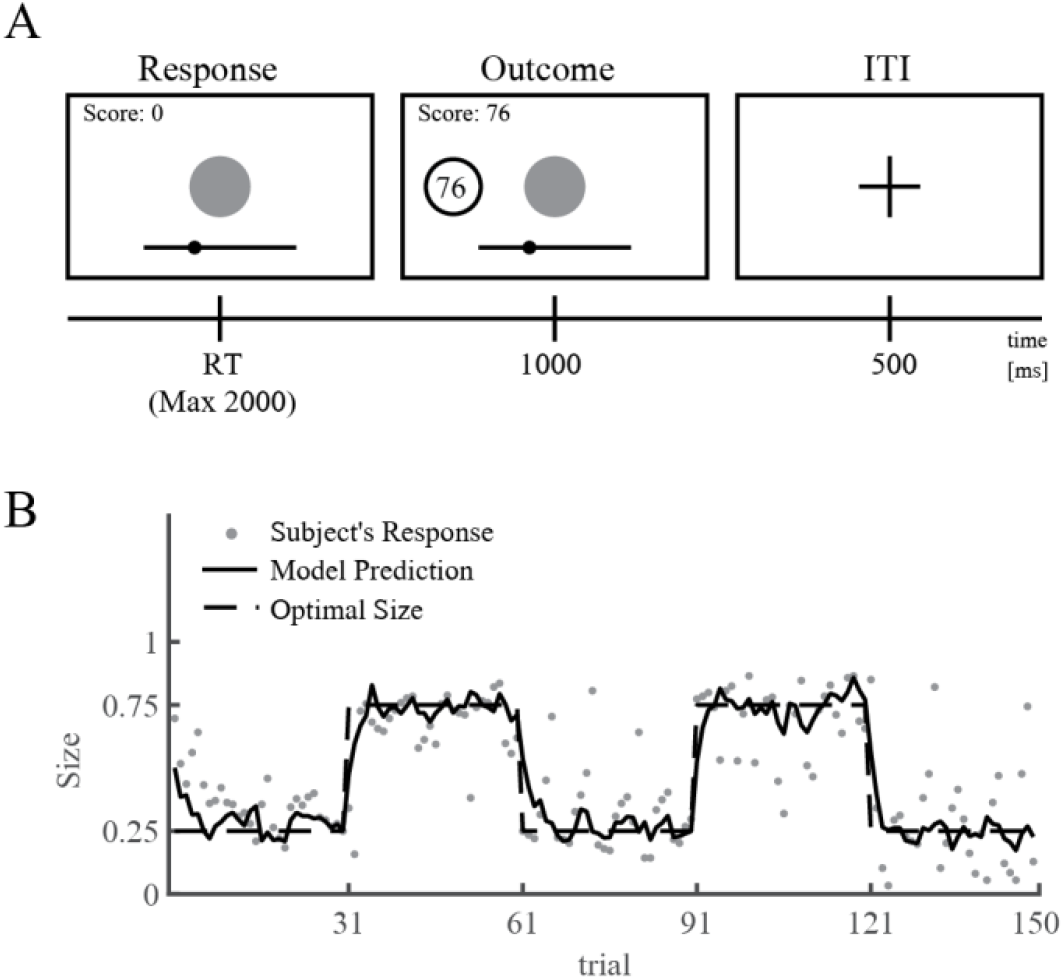
Continuous-score Probabilistic Reversal Learning Test (CPRLT). Panel A shows the structure of the task. The subject should select a size that considers to give a higher reward, and then he sees a reward between -100 to 100. Panel B shows a sample data and the fitted Rescorla-Wagner model to the data.

The best size to choose changes during the test. For the first 30 trials, small sizes are better. Then for the next 30 trials, large sizes are better. This switch happens every 30 trials (at trials 31, 61, 91, and 121). The score is calculated based on the difference of the participant response and a hidden size. In small size trials, the hidden size is drawn from a normal distribution with mean=0.25 and SD=1/12; In large size trials, the hidden size is drawn from a normal distribution with mean=0.75 and SD=1/12. Based on this, we also defined an “optimal size”, which is the size that gives the highest score if selected repeatedly: 0.25 for small size trials, and 0.75 for large size trials.

Before the main test, participants do 10 practice trials (the scores are random, and the first 3 trials have no time limit).

To calculate performance, we first computed the error for each trial:

“Error = Chosen size – Optimal size”

Then we calculated the mean absolute error (MAE) for all trials, and also for trials after the first rule change (trials 31 to 150, called adjusted MAE ( MAE_adj)). Missed trials were left out. Higher scores show lower cognitive flexibility. The test also includes a simple computational model, based on Rescorla & Wagner )1972). The model assumes that the subject has an expectation about the best scoring value (V) in each trial. The difference between the hidden value (λ) and V, is the prediction error(δ):

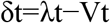

λt: Observed outcome at trial t; The hidden size in trial t

Vt: Current expected value; The subject’s belief about the best scoring size before trial t

The subject updates his/her expectation (V) according to the prediction error(δ), based on the following equation:

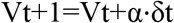

It gives each person a learning rate (α) between 0 and 1. A higher learning rate means quicker updates and better cognitive flexibility. We estimated α for each participant using maximum likelihood estimation.

## Analysis

To find the relation between age and cognitive flexibility scores, we used segmented regression models, instead of assuming a predefined shape (e.g. quadratic) for the relation. Segmented regression assumes that there is one or more point in which the slope of the linear relation changes. The model requires to pre-specify the number of break points. Therefore, based on the shape of the data, we fitted the model with one and two break points. Best model was selected by comparing the BIC of one break point model, two break point model, and simple linear model. We calculated 95% confidence interval for the peak age, peak cognitive flexibility score, and also for cognitive flexibility score along the age boundary, using bootstrap method. Participants with more than 10% missed trials or with a mean absolute error above 0.2 were classified as outliers and excluded from the final analysis.

## Results

The descriptive statistics and the details of the best segmented regression model were shown in table 1. Age varied widely, and the Kolmogorov–Smirnov test showed that age was not normally distributed (p = 0.5). The other variables were normal. For all scores, the model with 1 breakpoint either had the lowest BIC, or the difference of the BIC of the model with 1 breakpoint and 2 breakpoints was less than 2, and therefore we selected the simpler model. Figure 3 shows the visualization of the segmented regression models for every score.

**Table 1.**
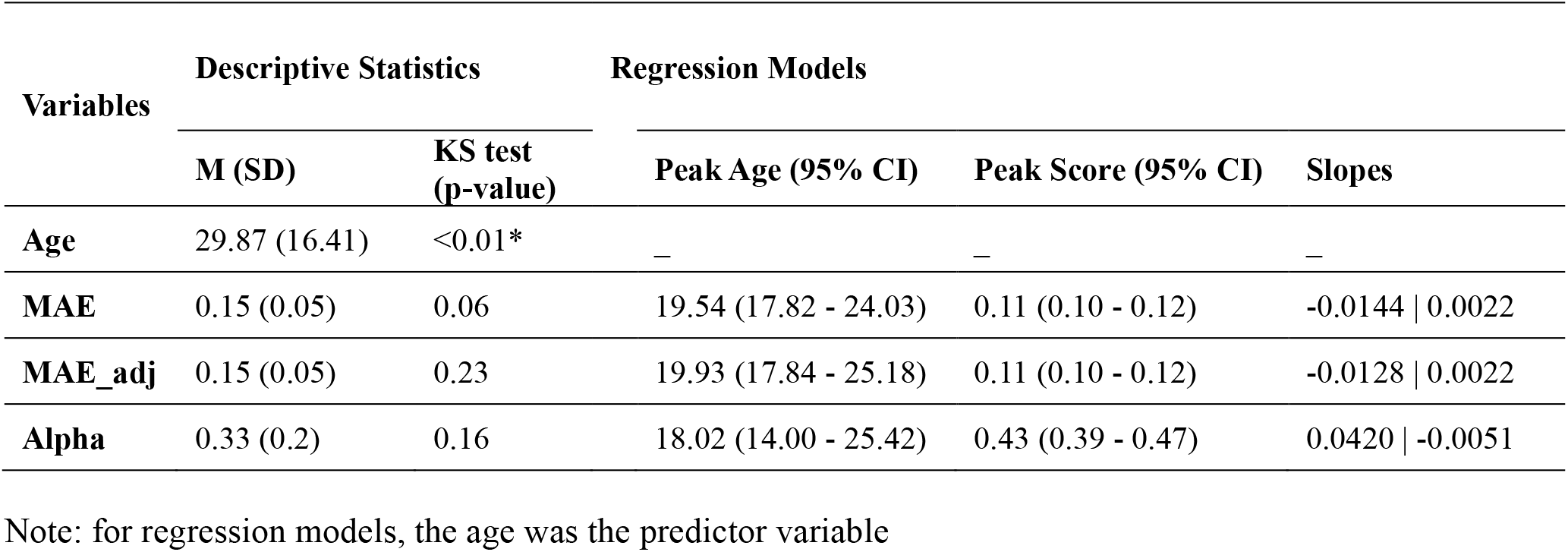
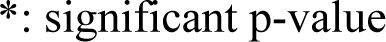
Descriptive statistics and regression model results.

| Variables | Descriptive Statistics |  | Regression Models |  |  |
| --- | --- | --- | --- | --- | --- |
|  | M (SD) | KS test (p-value) | Peak Age (95% CI) | Peak Score (95% CI) | Slopes |
| Age | 29.87 (16.41) | <0.01* | – | – | – |
| MAE | 0.15 (0.05) | 0.06 | 19.54 (17.82 - 24.03) | 0.11 (0.10 - 0.12) | -0.0144 \| 0.0022 |
| MAE_adj | 0.15 (0.05) | 0.23 | 19.93 (17.84 - 25.18) | 0.11 (0.10 - 0.12) | -0.0128 \| 0.0022 |
| Alpha | 0.33 (0.2) | 0.16 | 18.02 (14.00 - 25.42) | 0.43 (0.39 - 0.47) | 0.0420 \| -0.0051 |

Note: for regression models, the age was the predictor variable
\*: significant p-value

We found a single breakpoint for all three scores. For MAE, the breakpoint occurred at age 19.54 (95% CI [17.82, 24.03], Figure 2), with a peak error of 0.11 (95% CI [0.10, 0.12]). Before the breakpoint, error decreased by 0.0144 per year, indicating rapid improvement. After the breakpoint, error increased by 0.0022 per year, indicating a slow decline. MAE_adj showed a similar pattern, with a breakpoint at age 19.93 (95% CI [17.84, 25.18]) and a peak error of 0.11 (95% CI [0.10, 0.12]), with slopes of -0.0128 and +0.0022. For alpha (learning rate), the breakpoint occurred earlier, at age 18.02 (95% CI [14.00, 25.42], Figure 4), with a peak value of 0.43 (95% CI [0.39, 0.47]). Before the breakpoint, alpha increased by 0.0420 per year; after the breakpoint, it decreased by 0.0051 per year.

**Figure 2.**
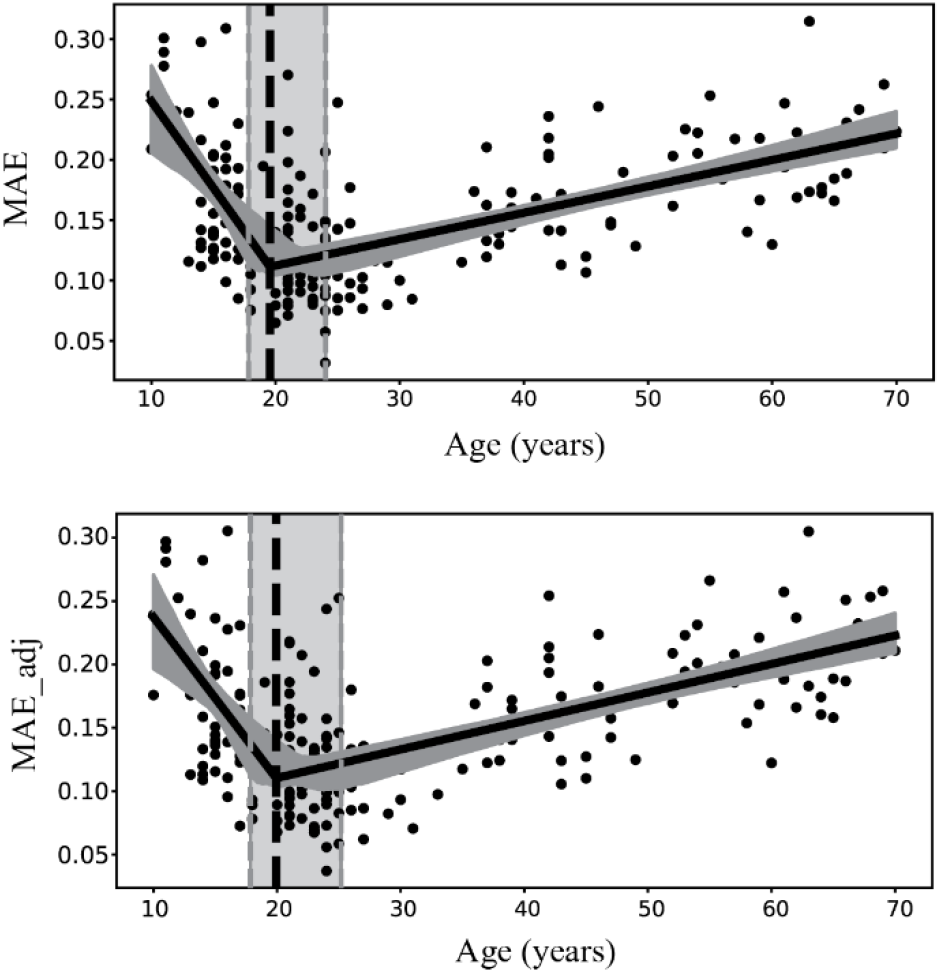
Divergent lifespan trajectories of rule-based and reward-based cognitive flexibility. Panels show observed data (scatterplots) and fitted segmented regression models (solid lines) with 95% CIs (shaded ribbons) for three CPRLT-derived scores across 221 healthy adults (aged 18–71).(Top panel) Mean Absolute Error (MAE) for rule-based learning. (Bottom panel) Adjusted Mean Absolute Error (MAE_adj) for rule-based learning. Vertical dashed lines and shaded bands indicate the estimated peak age and its 95% CI for each score, revealing that rule-based learning peaks later (∼age 20) than reward-based learning (∼age 18).

**Figure 3.**
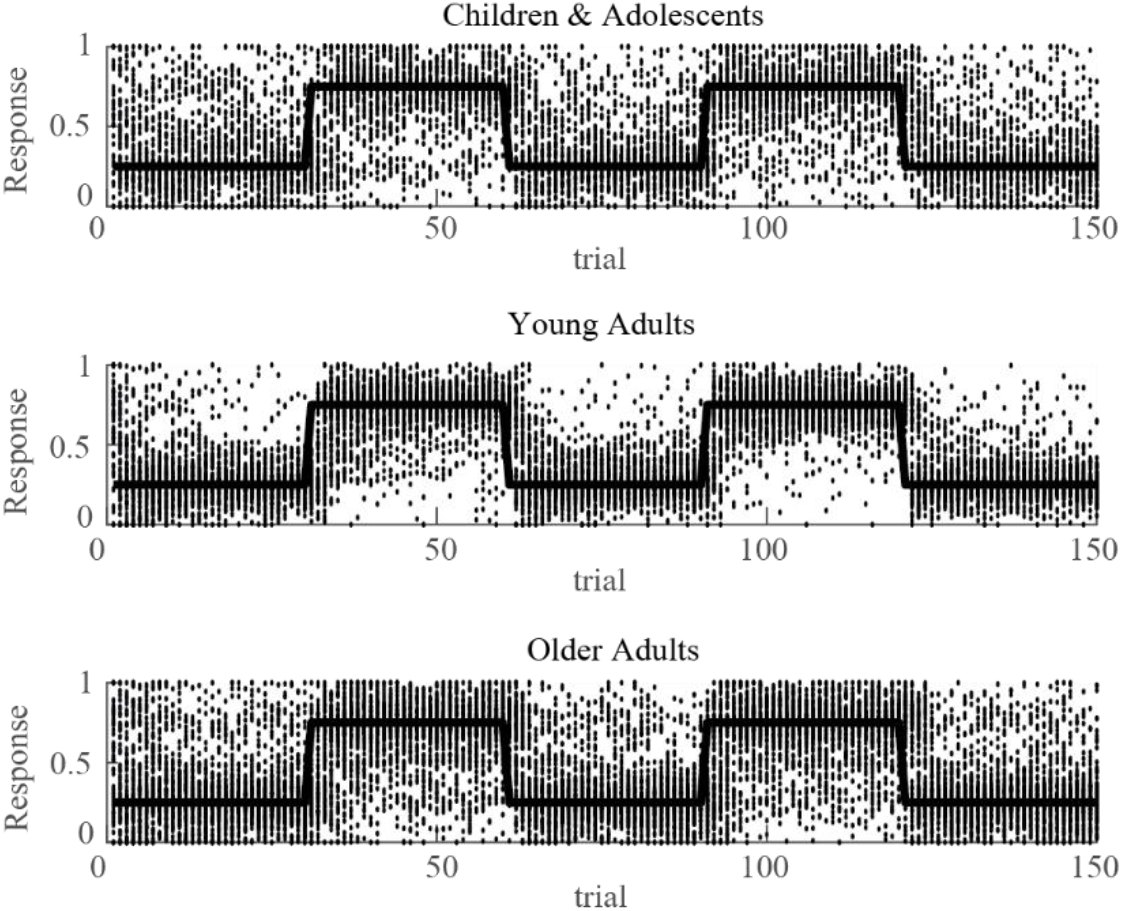
Overall plot for subject responses in CPRLT. We plotted selected sizes along trials for three groups of participants: Children and adolescents (age <18), Young adults (18<=age<=24), Older adults (age>24). Comparing the overall performance of these three groups, it can be seen that the young adults selected sizes closer to the optimal size, and adapted their response to change of the rule better than the other groups.

**Figure 4.**
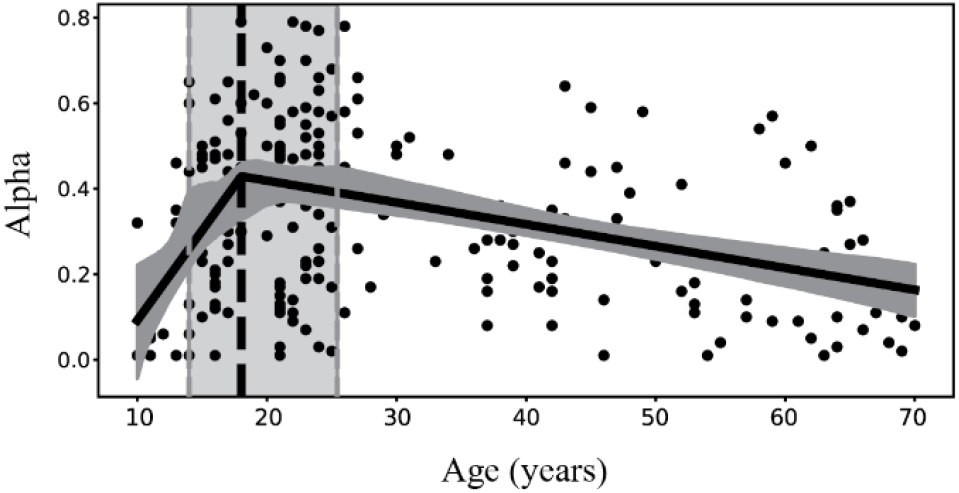
Divergent lifespan trajectories of learning rate (alpha) from the Rescorla-Wagner model for reward-based learning. Vertical dashed lines and shaded bands indicate the estimated peak age and its 95% CI for each score, revealing that rule-based learning peaks later (∼age 20) than reward-based learning (∼age 18) . Rule-based and reward-based cognitive flexibility. Panels show observed data (scatterplots) and fitted segmented regression models (solid lines) with 95% CIs (shaded ribbons) for three CPRLT-derived scores across 221 healthy adults (aged 18–71).

## Discussion

The results showed a rapid improvement in cognitive flexibility and reward-based learning from childhood to young adulthood, followed by a slower decline to old ages. The MAE and MAE-adj scores, which indicate rule-based learning, reach their peak at approximately 20 years of age, while learning rate (alpha), which indicates reward based free learning, reach its peak at approximately 18 years of age. This suggests that reward based free learning develops earlier than rule-based learning. This pattern is consistent with studies showing that prefrontal regions supporting rule-based behavior mature later than striatal circuits involved in reward processing (Tong et al., 2024).

Why would reward-based learning peak earlier? One possibility is brain development. Reward circuits, including the ventral striatum and orbitofrontal cortex, mature earlier in adolescence. Rule-based learning depends more on the lateral prefrontal cortex, which matures later into early adulthood (Kupis & Uddin, 2023). Our data fit that picture. The earlier peak for alpha and the later peak for MAE match what we know about these neural systems.

After the peak, white matter integrity and cortical thickness gradually decrease. A recent study showed that reduced white matter integrity in fronto-cerebellar tracts is associated with poorer executive function in older, but not younger, adults (Kraft et al., 2026). These neural changes don’t happen suddenly. Our behavioral data are consistent with that gradual shift. Other work has reported similar age-related declines in executive tasks that require flexible rule switching, particularly after age 30 (Cepeda et al., 2001; Huizinga et al., 2006).

Our task measured error on a continuous scale. Binary tasks would have missed the difference between these two peak ages. Continuous measures in working memory research have shown similar advantages (Bays et al., 2024; Esfahan et al., 2025). We show the same logic works for cognitive flexibility.

These findings have practical implications. Understanding that rule-based and reward-based learning peak at different ages can help design age-appropriate educational and training programs. For example, interventions targeting rule-based flexibility might be more effective in early twenties, while reward-based training could start earlier. Future longitudinal studies are needed to confirm whether the same individuals show this developmental sequence.

Limitations. Our data are cross sectional. A longitudinal study would be stronger, tracking the same individuals over time (Cole, 2024) argues that brain network dynamics change within individuals, so within subject data are needed. Also, we used only one flexibility task – probabilistic reversal learning. Other forms (task switching, attentional shifting) might age differently. Future studies should test multiple tasks.

